# Emergence of the Zika virus Asian lineage in Angola

**DOI:** 10.1101/520437

**Authors:** Sarah C. Hill, Jocelyne Vasconcelos, Zoraima Neto, Domingos Jandondo, Líbia Zé-Zé, Renato Santana Aguiar, Joilson Xavier, Julien Thézé, Marinela Mirandela, Ana Luísa Micolo Cândido, Filipa Vaz, Cruz dos Santos Sebastião, Chieh-Hsi Wu, Moritz Kraemer, Adriana Melo, Bruno L.F. Schamber-Reis, Girlene S. de Azevedo, Amilcar Tanuri, Luiza M. Higa, Carina Clemente, Sara Pereira da Silva, Darlan da Silva Candido, Ingra M. Claro, Nurse Domingos Quibuco, Cristóvão Domingos, Bárbara Pocongo, Alexander G. Watts, Kamran Khan, Luiz Carlos Junior Alcantara, Ester C. Sabino, Eve Lackritz, Oliver G. Pybus, Maria-João Alves, Joana Afonso, Nuno R. Faria

## Abstract

**Research In Context:** *Evidence before this study:* We searched PubMed without language restrictions using the keywords ‘Zika’ and ‘Africa’ for papers published to October 2018. We also checked available ‘Situation Report’ publications from WHO for evidence of Zika virus (ZIKV) or congenital Zika disease in Africa. ZIKV African lineage has been detected within Africa since the mid 20^th^ century, yet evidence for spread of the ZIKV Asian lineage within Africa is limited. Two countries in Africa (Cabo Verde and Angola) have reported ZIKV cases that are believed to be caused by a newly introduced Asian lineage virus. Sequence data are critical for confirming and understanding the spread of ZIKV Asian lineage within Africa, but these data are currently limited to a single 193bp fragment of the ZIKV NS1 gene from Angola. In addition, whilst epidemiological data on ZIKV and suspected microcephaly cases have been reported in detail from Cabo Verde, data from Angola are extremely limited.

*Added value of this study:* We provide a detailed report of detected ZIKV acute cases and suspected microcephaly cases in Angola. We sequence ZIKV genomes from three acutely infected cases. These represent the first three Asian lineage genomes available from Africa, one of which was acquired from a baby with confirmed microcephaly. Analysis of these sequences suggests that ZIKV may have been introduced to Angola between July 2015 and June 2016, after which it likely circulated for at least one year. This provides the first genetic confirmation of autochthonous ZIKV Asian lineage transmission within Africa. We suggest that the virus was more likely introduced to Angola directly from Brazil, rather than from Cabo Verde. Our analyses from Angola, only the second African country to report presence of the Asian virus lineage, therefore improve our understanding of the extent and clinical impact of ZIKV Asian lineage in the continent.

*Implications of all the available evidence:* The circulation of ZIKV Asian lineage within parts of sub-Saharan Africa is concerning given the potential for continued viral spread across much of the continent. Available evidence suggests that ZIKV has circulated and caused cases of microcephaly in Cabo Verde and in Angola. Detecting additional ZIKV transmission using only clinical data on suspected microcephaly or clusters of mild illness may be challenging in countries where systems for reporting birth defects are limited and infectious disease burden is high. Further spread of the ZIKV Asian lineage would likely not be detected unless molecular surveillance systems for ZIKV are implemented to routinely monitor ZIKV transmission in Africa. Implementation of such a surveillance system is especially important in countries that are linked by high human mobility to areas that have experienced recent or ongoing outbreaks of ZIKV.

**Abstract:** *Background:* Zika virus (ZIKV) infections and suspected microcephaly cases have been recently reported in Angola, but no data are available on the origins, epidemiology, and diversity of the virus.

*Methods:* Serum samples from 54 suspected ZIKV cases, 76 suspected microcephaly cases, and 24 mothers of infants with suspected microcephaly were received by the Angolan Ministry of Health. Computed tomographic brain imaging and serological assays (PRNT) were conducted on one microcephalic infant. All sera were tested for ZIKV by RT-qPCR. 349 samples from HIV+ patients and 336 samples from patients suspected of chikungunya virus or dengue virus infection were also tested. Portable sequencing was used to generate Angolan ZIKV genome sequences, including from a ZIKV+ neonate with microcephaly born in Portugal to an Angolan resident. Genetic and mobility data were analysed to investigate the date of introduction and geographic origin of ZIKV in Angola.

*Findings:* Four autochthonous cases were ZIKV positive via RT-qPCR, with all positive samples collected between December 2016 and June 2017. Viral genomes were generated for two of these cases, and from the neonate with microcephaly identified in Portugal. Genetic analyses and other data indicate that ZIKV was introduced to Angola from Brazil between July 2015 and June 2016. This introduction likely initiated local ZIKV circulation in Angola that continued until June 2017. The scanned microcephaly case showed brain abnormalities consistent with congenital Zika syndrome and serological evidence for maternal ZIKV infection.

*Interpretation:* Our analyses confirm the autochthonous transmission of the ZIKV Asian lineage in continental Africa. Conducting ZIKV surveillance throughout Africa is critical in the light of presented evidence for autochthonous ZIKV transmission in Angola, and associated microcephaly cases.

*Funding:* Royal Society, Wellcome Trust, CNPq, CAPES, ERC, Oxford Martin School, Global Challenges Research Fund, Africa Oxford, and John Fell Fund.

## Background

Zika virus (ZIKV) is an RNA virus of the *Flavivirus* genus that is primarily transmitted by *Aedes sp*. mosquitos. ZIKV is classified into two distinct lineages, the African and the Asian genotypes. Serological studies suggest that ZIKV may be widespread across Africa,^1^ yet the interpretation of serological assays is challenging due to extensive cross-reactivity among related flaviviruses.^2^ Prior to 2007, ZIKV was identified in only 14 human cases in Africa and Asia^3^ and infection was thought to cause only mild symptoms, including fever, headache and rash.^1^ However, since 2013, the Asian genotype of ZIKV spread to locations in the Pacific and the Americas, resulting in > 800,000 suspected and confirmed cases of Zika virus disease.^4^ The discovery that ZIKV infection during pregnancy can cause severe birth defects and other adverse outcomes,^2^ led to a research response that, to date, has been focused largely on the Americas.

Hundreds of millions of people in sub-Saharan Africa live in areas thought to be suitable for ZIKV transmission.^5^ Despite evidence of past transmission in Africa of the African genotype of ZIKV,^3^ there is a lack of data on recent ZIKV transmission from the continent. Since 2015, three African countries (Guinea-Bissau, Cabo Verde and Angola) have reported suspected human cases of ZIKV and clusters of suspected microcephaly cases.^6–9^ Understanding these ZIKV outbreaks is critical for safeguarding public health in Africa and elsewhere.

Obtaining accurate ZIKV surveillance data is challenging because most cases are asymptomatic, symptoms are mild and non-specific, and infections are frequently misclassified.^2^ In the absence of complete surveillance data, virus genome sequences have proven important for investigating ZIKV transmission.^10^ The current lack of ZIKV genomes from Africa hinders our understanding of the re-emergence of ZIKV in the continent.

Here, we provide the first comprehensive study of the recent ZIKV outbreak in Angola. We report and analyse confirmed ZIKV infected cases and suspected microcephaly cases. Computed tomographic (CT) imaging of one infant confirms the presence of brain abnormalities consistent with congenital Zika syndrome. The mother of this infant was serologically positive for ZIKV infection. We tested available serum samples for ZIKV and used portable sequencing technology to generate three Asian lineage ZIKV genome sequences from individuals infected in Angola, including one individual born with microcephaly.^11^ Phylogenetic analyses reconstruct the timings and geographic origin of the Angolan ZIKV outbreak. We corroborate our conclusions using multiple data sources, including the climatic suitability of the *Aedes* mosquito vector, international data on ZIKV incidence and airline passenger numbers. Our study is the first detailed analysis of the introduction to and circulation of ZIKV Asian lineage in Africa.

## Materials and Methods

### Zika virus surveillance and testing

The Ministry of Health of Angola conducted diagnostic testing of suspected cases of acute ZIKV infection at the Instituto Nacional de Investigação em Saúde (INIS) since late December 2016. Between late December 2016 and November 2018, 54 serum samples were collected from patients following clinical examination. Viral RNA was extracted from these samples using QiaAmp Viral RNA Mini Kits. RT-qPCR testing for the presence of ZIKV, chikungunya virus (CHIKV) and dengue virus (DENV) RNA was conducted using CDC Trioplex kits^12^ or Bio-Manguinhos (Fiocruz) ZDC RT-qPCR kits on an Applied Biosystems 7500 Fast machine.

On 1^st^ January 2018, it became mandatory for health providers in Angola to notify the Angolan Direcção Nacional de Saúde Pública (DNSP) of infants with suspected microcephaly (additional **Supplementary Materials and Methods** for case identification details). In total, 76 samples were received from infants between January 2017 and November 2018, with a median time of sample collection of 24 days after birth (mean 65 days, range 0-315 days) (**Figure S1**). Serum samples from 24 mothers who gave birth to infants with suspected microcephaly were also collected. All sera were tested by PCR as described above. Serological testing for toxoplasmosis, rubella and cytomegalovirus (ToRC pathogens) was conducted on eight serum samples from infants with suspected microcephaly using the ViDAS ToRC panel.

An additional 685 serum samples were investigated for ZIKV. Approximately half (n=336) were collected between January and October 2018 from patients with suspected DENV or CHIKV acute infections. These samples were PCR tested for DENV, CHIKV and ZIKV as part of routine diagnosis. The remaining 349 samples were collected between April and November 2017 by the Instituto Nacional de Luta Contra SIDA as part of a separate study investigating antiviral drug resistance among HIV+ patients in Luanda. These samples were retrospectively tested for ZIKV only, to improve the chance of detecting ZIKV cases.

Patients provided written consent forms agreeing that samples could be used for research purposes. Residual samples from Angola were used for viral genomic sequencing with the approval of the National Ethical Committee, Ministry of Health, Angola.

### Brain imaging of Angolan infant with microcephaly

CT imaging was conducted on one female child who was born in early August 2017 with suspected microcephaly. The child was born in Angola but diagnosed later at the Microcephaly Reference Centre IPESQ (Campina Grande, Brazil), following travel from Angola to Brazil in early November 2018. Plasma samples were taken from both the mother and child on 30^th^ November 2018 and tested for ZIKV and DENV infection using EuroImmun IgG ELISAs. Plaque reduction neutralisation tests (PRNT) were performed using standard protocols (**Supplementary Materials and Methods**) to quantitate neutralising antibodies against ZIKV for both the mother and the child. CT scans were conducted to assess whether the child’s neurological damage was consistent with congenital Zika syndrome. CT imaging was performed with a 64-section CT scanner (Philips Brilliance).

### ZIKV genome sequencing

Sequencing of the ZIKV genome coding regions was attempted using an Oxford Nanopore (ONT) MinION device following previously published protocols.^13^ Virus genome sequencing of RT-qPCR+ samples was attempted at INIS (Luanda, Angola). Generation of consensus sequences was undertaken using a previously published and validated bioinformatics pipeline (**Supplementary Materials and Methods**).^13^

Whilst investigations in Angola were underway, ZIKV RNA was detected in the urine of a neonate with microcephaly born in Portugal in November 2017 to a resident of Angola. Details of this case, a short (193 nt) fragment of the ZIKV NS1 gene from the neonate, and ethical permissions are described elsewhere.^11^ Virus genome sequencing of that strain was attempted using the methods described above, at the Instituto Nacional de Saúde Doutor Ricardo Jorge, Portugal.

### Phylogenetic analyses

The new ZIKV genomic sequences were aligned with 390 other ZIKV genomes publicly available from GenBank using Muscle.^14^ Maximum likelihood phylogenies were estimated using RAxML v8.2.11,^15^ and molecular clock phylogenies were generated in BEAST v1.10.3 ^16^ (see **Supplementary Materials and Methods** for further details).

### Origins of Angolan ZIKV outbreak

To corroborate the results of our genomic analyses, we investigated countries that could have exported ZIKV to Angola using additional data sources. Specifically, two factors were considered here as contributing to a high risk of ZIKV introduction to Angola: (i) countries with a high local incidence of ZIKV, and (ii) countries with high numbers of air passengers travelling to Angola. The monthly number of passengers to Angola from countries reporting ZIKV was estimated using worldwide ticket sales data from the International Air Transport Association during 2015-2017.^17^ The average incidence rate per person per week in each country was estimated from surveillance data on the number of suspected and confirmed cases of ZIKV per epidemiological week.^7,18^ See **Supplementary Materials and Methods** for details.

## Results

### Confirmed autochthonous ZIKV transmission in Angola

We collated evidence of ZIKV in Angolan residents or in travellers returning from Angola (**Figure 1**). ZIKV infection was first detected by seroneutralisation assay in a traveller returning from Angola in September 2016.^19^ In late December 2016, RT-qPCR testing of suspected acute ZIKV infections began at INIS. Between December 2016 and June 2017, four ZIKV cases were confirmed locally by Trioplex RT-qPCR. Three of these cases were identified among 54 symptomatic patients with suspected acute ZIKV infections. The remaining case (from June 2017) was detected through retrospective screening of an unrelated archive of 349 HIV+ samples obtained during April to November 2017. No additional ZIKV cases were detected among 336 samples collected from patients with suspected DENV or CHIKV infections during 2018. None of the ZIKV-positive samples were positive for DENV or CHIKV. Of the 51 ZIKV-negative samples from suspected ZIKV cases, three were positive for DENV-2, and seven were positive for CHIKV.

**Figure 1.**
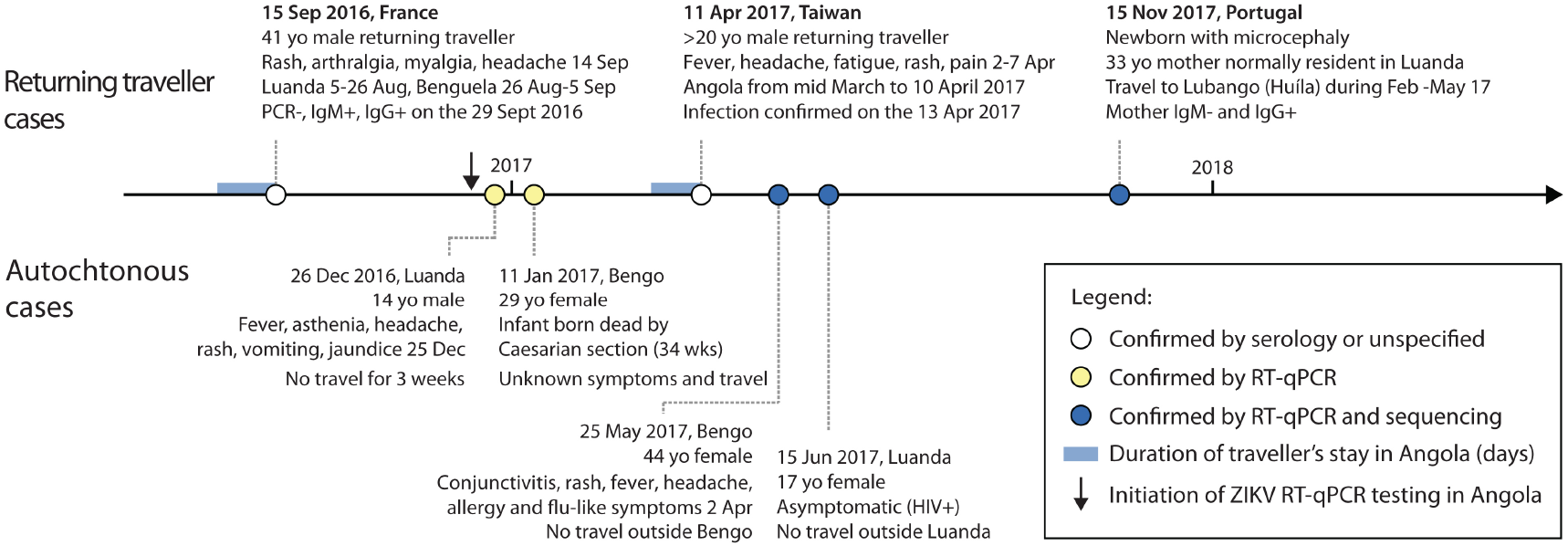
Confirmed Angola-associated ZIKV cases. Patient data and symptoms are reported where known. Travellers returning from Angola are shown above the line and local cases are shown below.

The presence of ZIKV in Angola during 2016 and 2017 was also indicated by the detection of ZIKV in two travellers; one to Portugal^11^ and another returning to Taiwan^20^ (**Figure 1**). Whilst suspected cases were reported across Angola (**Figure 2**), all confirmed cases were residents of (or had travel history involving) Luanda or the neighbouring province of Bengo (**Figures 1 & 2**).

**Figure 2.**
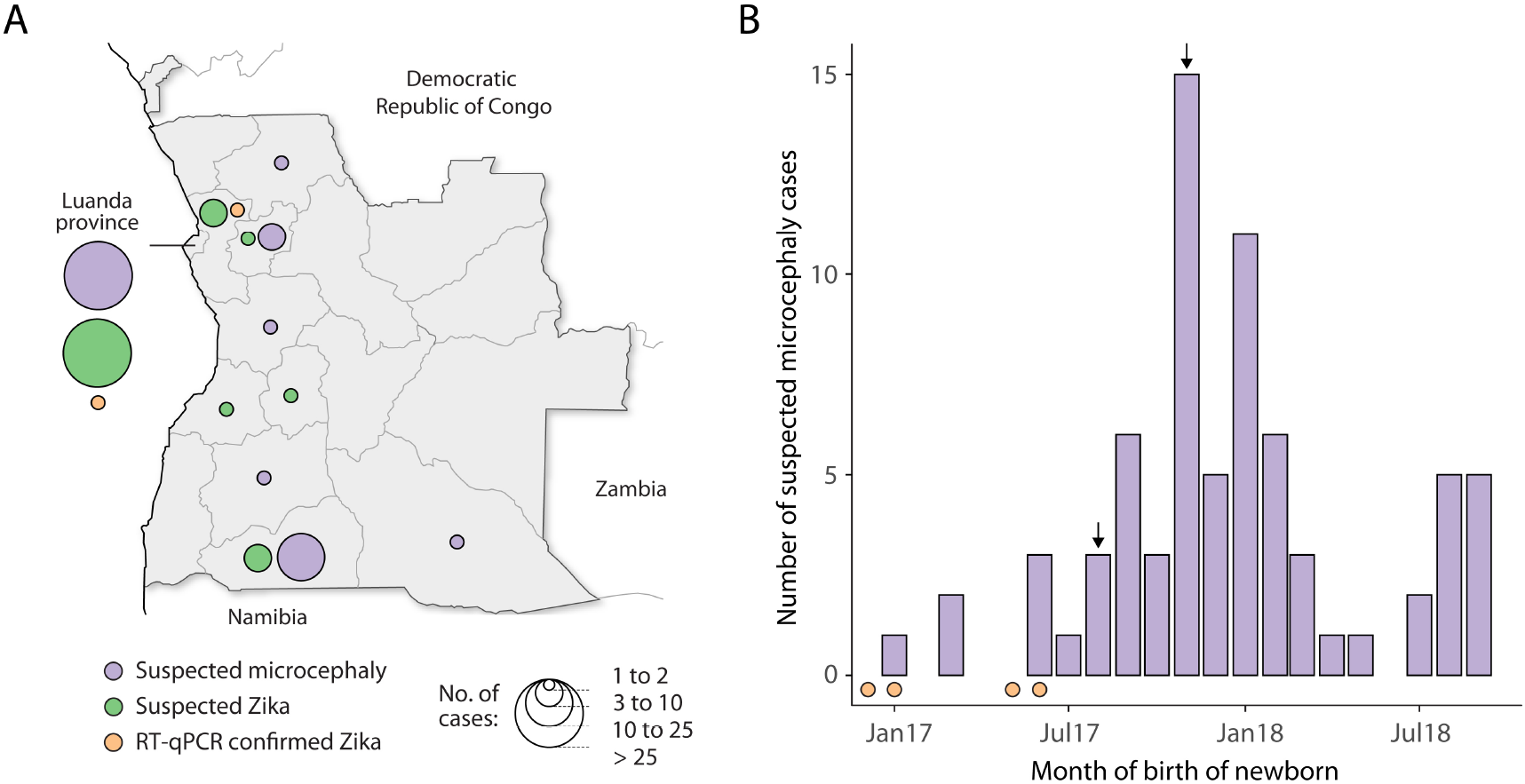
Suspected acute ZIKV cases and suspected microcephaly cases identified in Angola. A) Spatial distributions of suspected ZIKV cases and suspected microcephaly cases. Circle sizes indicate the number of cases per province. B) Time series of suspected microcephaly cases in Angola. Dates of birth, rather than report dates, are shown, so cases are included only if date of birth was recorded (96% of cases). Arrows mark the month of birth of two microcephaly cases that were independently identified and confirmed in Brazil (August 2017, confirmed here), and Portugal (November 2017 ^11^). Red dots on the horizontal axis indicate the sampling times of four PCR-confirmed ZIKV infections, identified in non-microcephalic patients in Angola.

### Microcephaly in Angola

Suspected microcephaly cases identified in Angola before November 2018 are shown in **Figure 2**. The number of neonates identified with suspected microcephaly peaks around November 2017, and subsequently declines (**Figure 2B**). The peak occurs several months after several PCR-confirmed ZIKV acute cases were detected in Angola (**Figure 2B**, red dots). However, the temporal and geographical distributions of suspected microcephaly and ZIKV cases are difficult to interpret with confidence because the total number of reported cases is low, the consistency of case detection and reporting are likely highly variable, and the intensity of surveillance efforts has changed over time (**Supplementary Materials and Methods**).

Of the 76 samples from infants with suspected microcephaly, none were positive for ZIKV by RT-qPCR. No maternal samples (n=24) tested PCR positive for ZIKV. This is not surprising, given that most infant samples were collected long after ZIKV is expected to be cleared from blood fluids^21^ (median time between birth and sample collection was 24 days, **Figure S1**), and because ZIKV infection may have occurred months prior to birth. Only 8 samples from infants with suspected microcephaly could be tested for infection with ToRC pathogens; none showed evidence of recent ToRC infection.

In addition to the suspected microcephaly cases reported to INIS, one Angolan microcephaly case was identified in Brazil. The 21 year-old mother of this child was a long-term resident of Moxico province in Angola, but travelled to Luanda during the second and third months of pregnancy and reports a rash during this visit (week 10 of pregnancy; January 2017). Confirmed ZIKV cases were identified in Luanda and Bengo province at this time (**Figure 1**). The neonate was delivered in early August 2017 in Angola, coincident with the rise in suspected microcephaly cases (**Figure 2**). CT scans of the child’s brain conducted at 15 months confirmed microcephaly through reduced cerebral volume, and showed abnormalities consistent with congenital Zika syndrome observed in Brazil and elsewhere^22^ (**Figure 3**). When tested serologically, the child was IgG negative for both DENV and ZIKV by ELISA, and weakly positive for neutralising antibody response by PRNT (PRNT90 titre = 40). This result does not exclude intrauterine ZIKV exposure, as the child was 15 months -old at testing (see **Materials and Methods**) and maternally acquired IgG antibodies are present for only 6-12 months after birth.^23^ The mother was strongly IgG positive for ZIKV by ELISA, with a weaker DENV IgG response. PRNT assays confirmed the strong neutralising antibody response in the mother, with a PRNT90 titre of 1280. This result strongly suggests that the mother had been previously infected with ZIKV.

**Figure 3.**
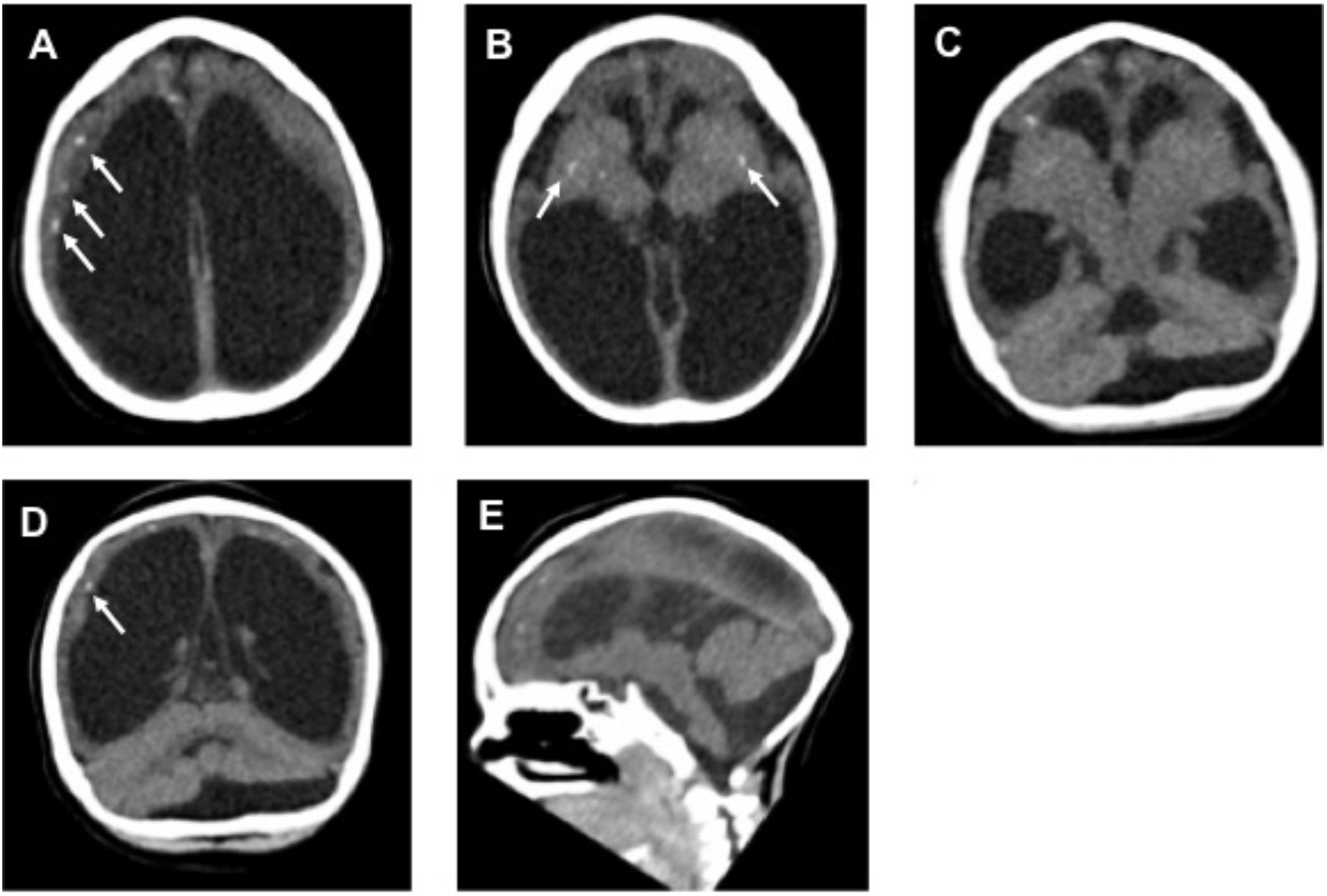
Brain CT image of an Angolan child with microcephaly. (A) Compensatory ventriculomegaly and calcification areas in the subcortical region (arrows). (B) Calcification in the basal ganglia (arrows). (C) Pachygyria. (D) Dysgenesis of the cerebellum and calcification in subcortical region (arrows). (E) Brainstem hypoplasia.

### Sequencing and genetic analysis of ZIKV

We sequenced three virus genomes from patients infected with ZIKV in Angola. **Table S1** provides sequencing statistics and GenBank accession numbers. The three Angolan ZIKV sequences differ from each other at 34 nucleotide sites. Three of these variable sites cause amino acid changes: Y135H (in sample Z3), Y3038H and D3344E (both in sample PoHuZV_634939, isolated from a microcephaly patient^11^). Neither of the two amino acid changes observed in strain PoHuZV_634939 are seen in either of the two most closely related ZIKV genomes obtained from infants born with microcephaly in Brazil (GenBank accession numbers KU729217, KU527068).

Maximum likelihood (ML) phylogenies indicate that all three ZIKV strains from Angola form a single, well-supported monophyletic clade within the ZIKV Asian genotype lineage currently circulating in the Americas (**Figure S3**; bootstrap score = 0·96). Thus the ZIKV outbreak in Angola likely resulted from a single successful introduction of the virus, followed by autochthonous transmission within Angola.

We used a Bayesian phylogenetic framework to investigate when and from where ZIKV was introduced to Angola. Our analyses indicate that ZIKV was likely introduced to Angola from Brazil (**Figure 4**), possibly from the southeast of Brazil, where the most closely related virus to the Angola ZIKV clade was sampled (accession number KY559016; posterior probability = 0·99). The estimated date of the most recent common ancestor of the ZIKV Angolan sequences is June 2016 (Bayesian 95% credible interval, January 2016-October 2016) (**Figure 4**). The date of divergence of the Angolan ZIKV cluster from the most closely related Brazilian ZIKV sequence is July 2015 (February 2015-February 2016) (**Figure 4**). These estimates therefore indicate that ZIKV was introduced to Angola from Brazil between July 2015 and June 2016. Thus ZIKV may have circulated in Angola for 3 to 14 months before being identified for the first time, via a traveller returning from Angola.^19^

**Figure 4.**
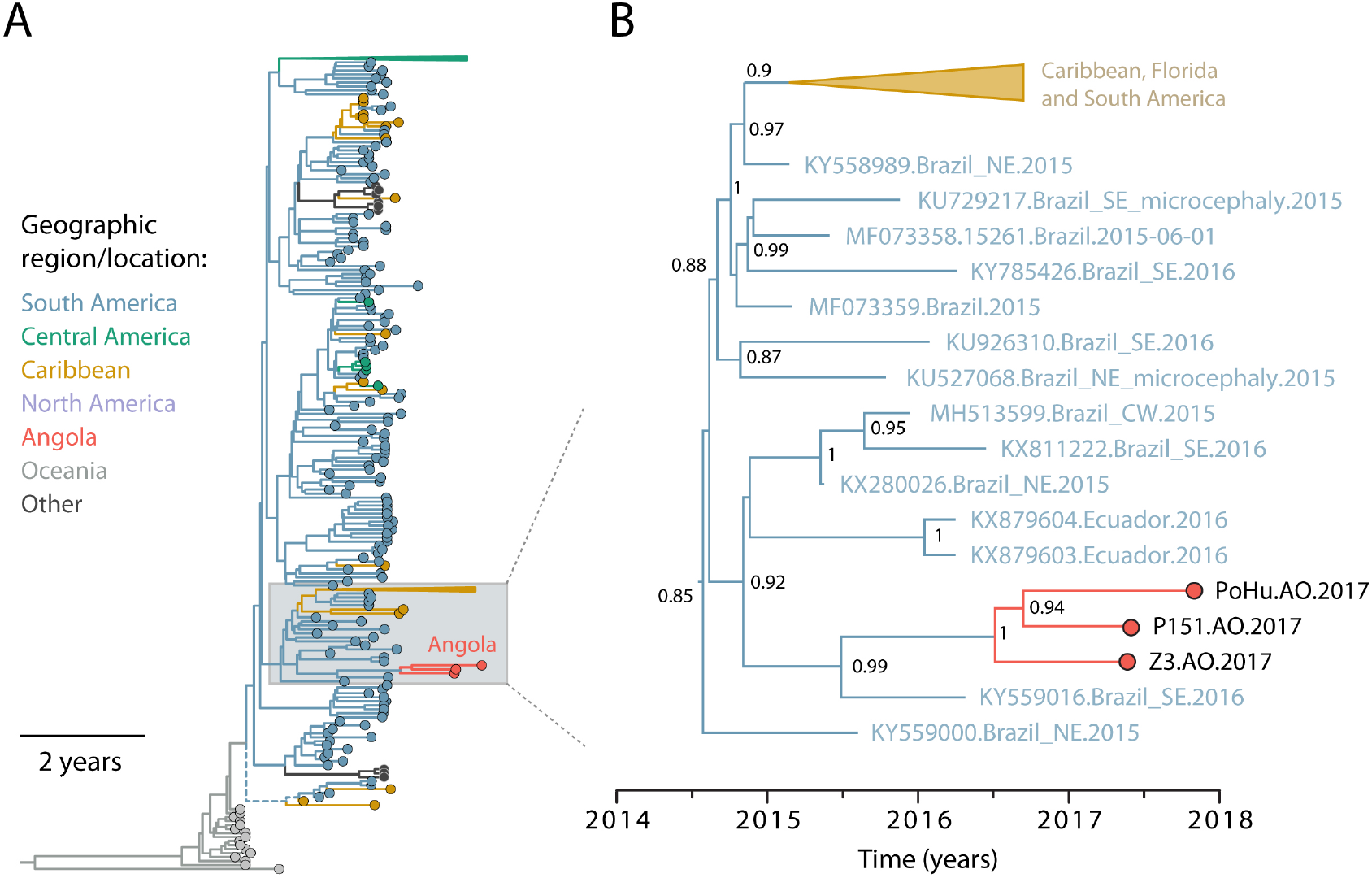
Phylogenetic analysis of the introduction of ZIKV to Angola. A) Maximum clade credibility phylogeny, estimated from complete and near-complete ZIKV genomes using a molecular clock phylogeographic approach. Branch colours indicate the most probable locations of ancestral lineages. Triangular clades represent larger groups of sequences that have been collapsed for visual clarity. B) Expansion of the clade in panel A containing the Angolan ZIKV (green) and closely related sequences from the Americas (red). Clade posterior probabilities are shown at well-supported nodes.

### Evaluation of the outbreak’s geographic origin

A ZIKV outbreak was reported in Cabo Verde during 2015-2016, from which no genetic sequence data are currently available.^7^ The absence of ZIKV genetic data from key locations, such as Cabo Verde (**Figure 5**), means that we cannot unambiguously infer the geographic origin of the Angolan ZIKV outbreak using phylogenetic analysis alone. We therefore analysed global ZIKV incidence and human mobility data to predict the likely source of ZIKV in Angola.

**Figure 5.**
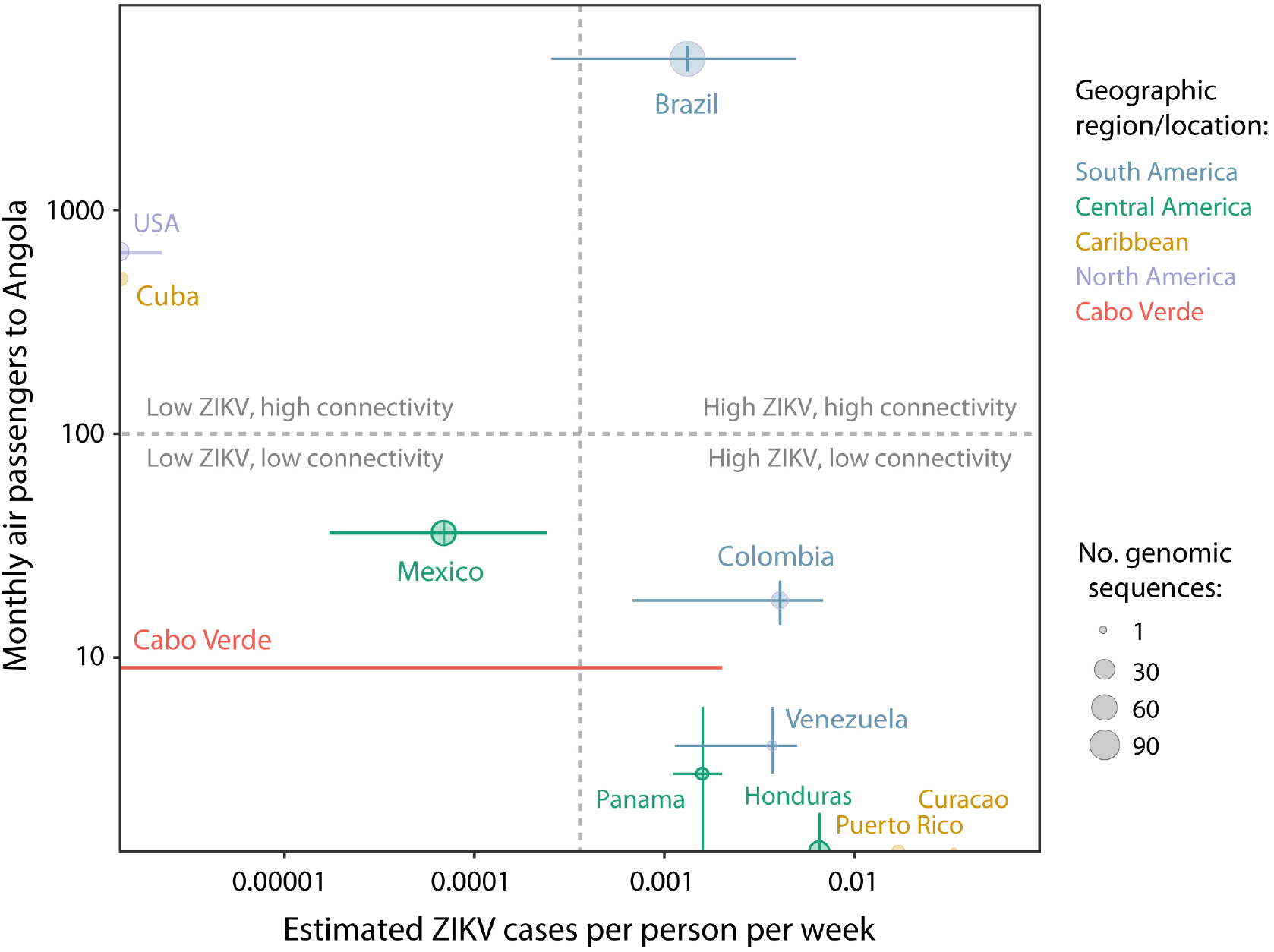
Factors affecting the likelihood of introduction of Asian lineage ZIKV into Angola. The vertical axis shows the median number of air passengers travelling to Angola per month during the most likely time for the most recent common ancestor of the Angolan ZIKV cluster (January 2016 – October 2016). Vertical bars show the 25% and 75% quartiles of monthly passenger numbers. The horizontal axis shows the median weekly number of suspected or confirmed ZIKV cases per person during the same period. Horizontal bars show the quartiles of ZIKV case numbers. Point diameters are proportional to the total number of genomic sequences currently available from each country. The 11 countries shown include all those with the with the seven highest median passenger numbers and the seven highest number of ZIKV cases per person. Colour indicate geographic region (red: Africa, green: Central America, purple: South America, blue: USA, orange: Caribbean).

We found that 80% of all air passengers who travelled to Angola from countries reporting ZIKV began their journey in Brazil, whilst only 0·15% originated from Cabo Verde. The number of passengers that travelled to Angola per month was therefore over 500 times higher from Brazil than from Cabo Verde (**Figure 5**). Crucially, Brazil is the only country that exhibits both high connectivity with Angola via air travel and a high average rate of ZIKV cases per person (**Figure 5**, top right). Further, the climatic suitability for Aedes-borne viral transmission was more closely synchronised between Angola and Brazil than Angola and Cabo Verde (**Figure S4**). Introduction of ZIKV to a new location is most likely to be successful when environmental suitability for the mosquito vector is high in both the source and destination locations. Thus, multiple lines of evidence (ZIKV incidence, mobility data, and virus phylogenetic analysis) support the hypothesis that ZIKV was introduced directly to Angola from Brazil.

## Discussion

Here, we characterise the first known outbreak of Asian lineage ZIKV in continental Africa. We tested 54 suspected cases of acute ZIKV infection and 76 suspected microcephaly cases sampled across Angola by RT-qPCR. Three suspected cases of acute ZIKV infection, all notified between December 2016 and May 2017, were ZIKV positive. A further ZIKV+ case was detected in a cohort of 349 HIV+ patients sampled in Luanda in June 2017. To understand the origins of the Angolan outbreak, we sequenced ZIKV genomes from two autochthonous cases and a confirmed microcephalic neonate, born in Portugal to a resident of Luanda. These represent the first ZIKV Asian lineage genomes reported from Africa, and the first genome from a ZIKV associated microcephaly case in Africa.

We analysed suspected microcephaly cases reported to INIS during 2017-2018. The fact that we found no ZIKV-positive cases among 76 suspected microcephaly cases does not indicate the virus was absent from this group, because the samples were collected long after birth, at which point ZIKV viremia is expected to have declined to undetectable levels in blood fluids.^21^ To maximise ZIKV detection sensitivity in suspected microcephaly cases, sera and urine samples for RT-qPCR should be collected as soon as possible after birth, and ideally within the first few days.^24^ Systems for the detection and reporting of birth defects are limited in sub-Saharan Africa, so the spatiotemporal distribution of suspected cases must be interpreted cautiously. In addition to the 76 suspected microcephaly cases identified in Angola, two cases of microcephaly acquired in Angola have been confirmed in other countries. Both cases (one of which is clinically reported here) have strong serological evidence for maternal ZIKV exposure and brain abnormalities consistent with congenital Zika syndrome.^11,22^ Further, both confirmed microcephaly cases were born a few months after the date of sampling of PCR-confirmed ZIKV infections in Angola. A time lag of ~5 months between ZIKV incidence and occurrence of microcephaly cases has been observed elsewhere.^25^ We show here that ZIKV detected in one these confirmed microcephaly cases^11^ belongs to a virus lineage originating from the Americas, and which likely circulated in Angola for at least 12 months. Continued identification of confirmed cases of acute ZIKV infection and ZIKV-associated microcephaly in Angola remains a priority to understand the clinical consequences of ZIKV in Africa.

Phylogenetic analysis shows that the three Angolan ZIKV genomes form a single clade whose common ancestor dates to June 2016 or earlier. Successful introductions of ZIKV to new regions often occur during seasons of peak climatic suitability for *Aedes-borne* transmission.^10,26^ Under this scenario, ZIKV would most likely have entered Angola between November 2015 and April 2016 (**Figure S4**). Detection of a single ZIKV outbreak clade in Angola is compatible with two scenarios, (i) a single successful introduction that initiated local ZIKV transmission in Angola, continuing until at least June 2017 or (ii) recurrent but later introduction to Angola of viruses belonging to a specific ZIKV lineage present in Brazil. The latter is much less likely, as it is improbable that three or more independent introductions of ZIKV to Angola belonged to just one of the many different lineages of ZIKV that co-circulated in Brazil.^10^ If the former scenario is correct, then ZIKV may have circulated undetected in Angola for at least 3 months before a case was first detected in a returning traveller,^19^ and at least 6 months before a local case was detected (**Figure 1**). Similar or longer periods of cryptic ZIKV transmission have been reported in the Americas^10^ and attributed in part to the difficulties in identifying clinical cases when infections are asymptomatic or mildly symptomatic.^27^ Retrospective screening of stored samples and identification of young children with suspected microcephaly, will help to determine the magnitude and duration of undetected ZIKV transmission in Angola since 2015.

We conclude that ZIKV was introduced to Angola from Brazil. This is consistent with previous predictions that, given the presence of ZIKV in Brazil, Angola is the African country most at-risk of importing the virus.^28^ Transmission of mosquito-borne viruses between these two countries was demonstrated previously by the spread of CHIKV East-Central-South African genotype (ECSA) from Angola to Brazil in June 2014.^29^ The introduction of ZIKV to Angola underscores the need to coordinate viral surveillance strategies across countries that share high human interconnectivity and similar vector-borne transmission potential, regardless of the transnational distance involved. Detecting ZIKV transmission using only clinical data on clusters of mild illness or suspected microcephaly would be extremely difficult in this setting. Molecular surveillance systems are therefore needed to monitor further ZIKV spread in Africa, including those countries potentially at risk of ZIKV introduction from Angola.^30^

The epidemiology and clinical significance of Asian and African ZIKV lineages in Africa is unclear. Widespread screening of pregnant women for STORCH pathogen infections, sensitive ZIKV antibody-based assays, and follow-up of children born with abnormalities would improve our understanding of the extent and causes of birth defects in Africa. Additional genomic surveillance of other mosquito-borne viruses, such as dengue and Japanese encephalitis virus, is required to inform effective intervention strategies to ameliorate disease burden in Angola. Previously, genomes of emerging viruses have been typically generated in a small number of highly resourced genomic centres, located in wealthy nations. The introduction of low cost, portable sequencing technology to laboratories in Africa heralds an era in which virus genome sequencing is widespread, real-time, and sustainable, with the potential to directly inform public health responses to future emerging outbreaks.

## Contributions

SCH performed laboratory work and analyses, and wrote the manuscript. JV, ZN, DJ, JX, MM, ALMC, FV, CSS, CC, SPS, DQ, CD, BP and IMC performed laboratory work on samples identified in Angola and epidemiological surveillance. LZZ and MJA identified and helped with sequencing of the Portuguese case. RSA assisted laboratory work in Angola, and performed investigations on the microcephalic infant identified in Brazil along with AM, BLFSR, AT, LMH and GSA. CHW helped with analyses. AGW, KK, MK and DSC assisted with analysis of global mobility and incidence data. JT and NRF conducted bioinformatics and assisted phylogenetic analyses. LCJA and ECS oversaw parts of the laboratory work. EL and OGP contributed to the overall design, provided interpretation of the results and commented on manuscript drafts. NRF and JA led the design and execution of the study, and oversaw all analyses and interpretation. All authors have seen and approved of the final text.

## Acknowledgements

N.R.F. is supported by a Sir Henry Dale Fellowship (204311/Z/16/Z), internal GCRF grant 005073, and John Fell Research Fund grant 005166. This research was supported by CNPq (440900/2016-6) and CAPES (88881.130757/2016-01). Travel was supported by Africa Oxford grants, AfiOx-48 (NRF) and AfiOx-60 (SCH). This work was supported by the European Research Council under FP7 grant agreement 614725-PATHPHYLODYN, and by the Oxford Martin School. Thanks to Dr. A.G.P Ferreira and Dr. P. Alvarez (Fundação Oswaldo Cruz (Fiocruz), Instituto de Tecnologia em Imunobiológicos Bio-Manguinhos, Rio de Janeiro, Brazil) for donating Bio-Manguinhos reagents. Thanks to Prof. J. Muñoz-Jordan and Dr. G. Santiago for donating RT-qPCR reagents.

## Data sharing

Sequence alignments, XMLs and tree files are available at GitHub (XXX to be completed prior to acceptance XXX). ZIKV genome sequences are available on GenBank with accession numbers XXX-XXX (pending).

