## Supplementary Materials and Methods for "Emergence of the Zika virus Asian lineage in Angola"

**Contents:**

Page 3: *Supplementary Materials and Methods.*

Page 5: *Supplementary Figure 1*. Number of days between birth and sample collected for suspected microcephaly cases.

Page 6: *Supplementary Figure 2*. Root to tip regression of sequence sampling date and genetic divergence and maximum likelihood phylogeny.

Page 7: *Supplementary Figure 3.* Maximum likelihood phylogeny of ZIKV

Page 8: *Supplementary Figure 4.* Seasonal climatic suitability for *Aedes aegypti*.

Page 9: *Supplementary Table 1.* Sequencing statistics for Angolan ZIKV genomes.

Page 10: *References.*

### Supplementary Materials and Methods

#### *ZIKV surveillance and diagnostic testing*

On 1st January 2018, it became mandatory for health providers in Angola to notify the Angolan Direção Nacional de Saúde Pública (DNSP) of infants with suspected microcephaly. Cases of suspected microcephaly were defined by the DNSP as newborns with cephalic perimeters at birth of <32 cm for males, or <31.5 cm for females, regardless of gestational age. This measure is equivalent to two standard deviations below the median cephalic perimeter expected for a *full term* birth.[1] Results of any additional clinical assessment and exact gestational age were not typically reported, so we stress that microcephaly is suspected but unconfirmed. When possible, sera were collected from infants born with suspected microcephaly and sent to INIS for serological and RT-qPCR testing for ZIKV. In total, 76 samples from infants were received for testing between January 2017 and 31st October 2018, with a median time of sample collection of 24 days after birth (mean 65 days, range 0-315 days) (**Figure S1**). This includes 17 samples collected during 2017, when case reporting of infants born with suspected microcephaly was not mandatory. Note that additional cases from which samples were *not* taken may have been reported to DNSP, but these cases that lack samples are not included here.

To improve the chance of detecting ZIKV cases, 685 additional serum samples were also tested by RT-qPCR. Samples from the Instituto Nacional de Luta Contra Sida were pooled in triplicate prior to RNA extraction and RT-qPCR, followed by individual testing of samples if a pool was positive. Testing of these, and other samples, is described in the **Main Text**.

#### *PRNT on microcephalic infant and mother*

Plaque reduction neutralisation tests (PRNT) were performed to quantitate neutralising antibodies against ZIKV. Briefly, plasma samples were heat-inactivated at 58°C. Two-fold dilutions of heat-inactivated plasma (ranging from 1:5 to 1:2,560) were incubated with 100 plaque forming units (PFU) of ZIKV (strain MR766) for 1 hour at 37°C. The virus-sera mixture was inoculated onto confluent monolayers of VERO cells. In addition, virus-only controls were included to determine the infectivity of the challenge virus. After 1 hour, inoculum was removed, and cells were overlaid with semisolid medium (1.25% carboxymethylcellulose in alpha-MEM supplemented with 1% fetal bovine serum) and then further incubated at 37°C for 5 days. Cells were fixed with 4% formaldehyde solution and stained with crystal violet dye solution for plaque visualisation. The plaque reduction neutralisation titre was defined as the highest serum dilution that results in 90% reduction (PRNT90) of infectivity when compared with the challenge virus.

#### *ZIKV sequencing and consensus sequence generation*

Sequencing of coding regions of the ZIKV genome was attempted using an Oxford Nanopore Technology (ONT) MinION device following previously published methods.[2] Briefly, cDNA was generated from viral RNA using random hexamers Protoscript II First Stand cDNA Synthesis kit or Superscript IV First Strand Synthesis System. Multiplex PCR with 42 cycles was used to generate overlapping amplicons that spanned the whole coding region of the ZIKV genome, according to previously published thermocycling conditions.[2] PCR products were purified using 1x Ampure XP beads, the concentration of DNA quantified using a Qubit Fluorometer, and samples were standardised by concentration. Library preparation was performed using 200-350 ng total mass of PCR product as input. Library preparation was performed using the ONT SQK-LSK108 ligation sequencing kit and NBD103 Native Barcoding Kit according to the manufacturer's instructions but with the changes detailed in [2] and https://www.protocols.io/view/one-pot-ligation-protocol-for-oxford-nanopore-librk9acz2e. The library was loaded onto FLO-MIN106 flow cells, and sequencing conducted without basecalling for 12-48 hours using MinKNOW 2 software. Negative controls that were extracted with the positive samples were also sequenced.

Raw sequencing data were processed according to established pipelines.[2] Raw reads were basecalled using Albacore (Oxford Nanopore Technologies), demultiplexed and adaptor-trimmed using Porechop, and mapped to a reference genome (Genbank accession number KJ776791) using bwa v 0.7.16a-r1181. Nanopolish software was used to identify variation from this reference genome. A consensus was generated for all genomic sites where coverage was at least >20X.

#### *Phylogenetic analysis*

The maximum likelihood tree was estimated using RAxML version 8.2.11 [3] under a general time reversible nucleotide substitution model, with gamma-distributed among-site rate variation and a proportion of invariant sites (GTR + G + I). Appropriate temporal signal for estimation of molecular clock phylogenies was assessed using TempEst [4] (**Figure S2**). Phylogenies calibrated in time units were estimated under a relaxed clock model and a codon-partition (SRD06) nucleotide substation model [5] for ZIKV using the MCMC approach implemented in BEAST v1.10.3.[6] Three independent MCMC chains of 250 million steps were computed, sampled every 25,000 steps. The first 10% of each run was discarded as burn-in, convergence of the runs was checked using Tracer 1.7.1,[7] and maximum clade credibility trees were constructed using TreeAnnotator.[6] Alignments, XMLs and tree files are available at GitHub (XXX to be completed prior to acceptance XXX).

#### *Estimation of the origins of Angolan ZIKV*

To further investigate the geographic origin of ZIKV in Angola, countries that could have exported the ZIKV lineage to Angola were identified using multiple data sources. Specifically, two factors were considered here as resulting in a high risk of introducing ZIKV to Angola: (i) countries with a high local incidence of ZIKV, and (ii), countries with high number of passengers travelling by air into Angola. Countries in which ZIKV Asian lineage confirmed cases were detected during 2015-2016 were identified from WHO reports.[8] For each of these countries, the number of passengers to Angola was estimated by analysing worldwide air ticket sales data from the International Air Transport Association (IATA) during 2015-2017.[9] These data included the full itineraries of travellers: their initial airport of embarkation, their final destination airport and, where applicable, connecting airports, but did not detail incomplete trips (e.g., due to missed flights).

Records of the number of suspected or confirmed cases of ZIKV from each country in the Americas that has reported ZIKV cases during the likely period of ZIKV introduction to Angola were used as a measure of local ZIKV incidence. Data from countries in Asia were not included as we show later that the Angolan ZIKV strain clearly belongs to the lineage of the Asian ZIKV genotype circulating in the Americas (**Figure 4**). Surveillance data for the Americas included suspected and confirmed cases of ZIKV per epidemiological week reported to PAHO.[10] Equivalent data for Cabo Verde, the only other country in Africa reporting the ZIKV Asian lineage, were taken from elsewhere.[11] Only suspected and confirmed cases that occurred around the time of the most recent common ancestor of ZIKV strains in Angola were considered (as estimated using our molecular clock analysis). To provide a crude estimate of infection risk per person, here we scale the number of reported cases by the population size in each country in 2015.[12]

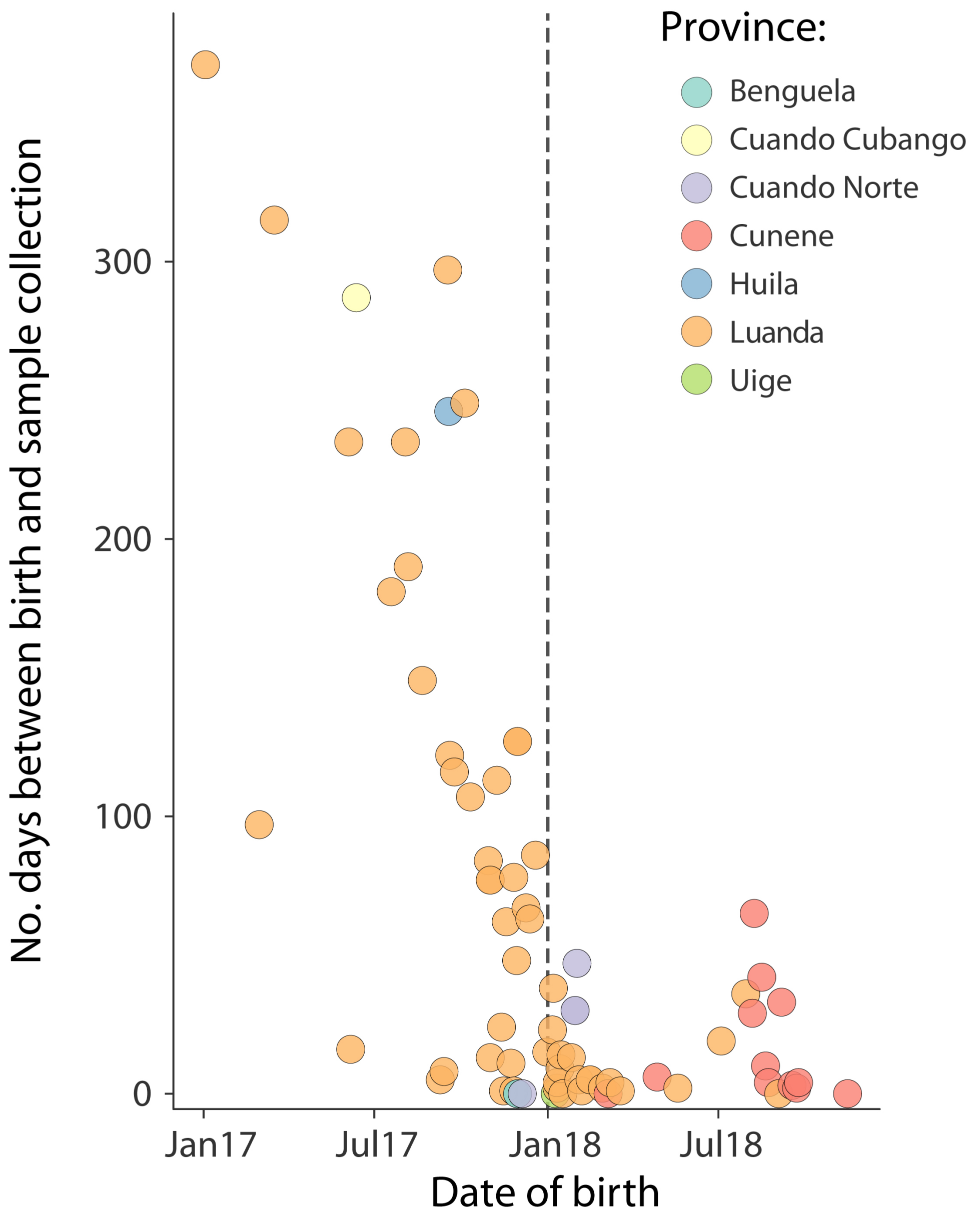


**Supplementary Figure 1. Number of days between birth and sample collected for suspected microcephaly cases.** Colours indicate province of sampling. Dashed vertical line indicates implementation of mandatory case reporting of babies with suspected microcephaly in Angola. Most cases identified prior to this time were identified retrospectively.

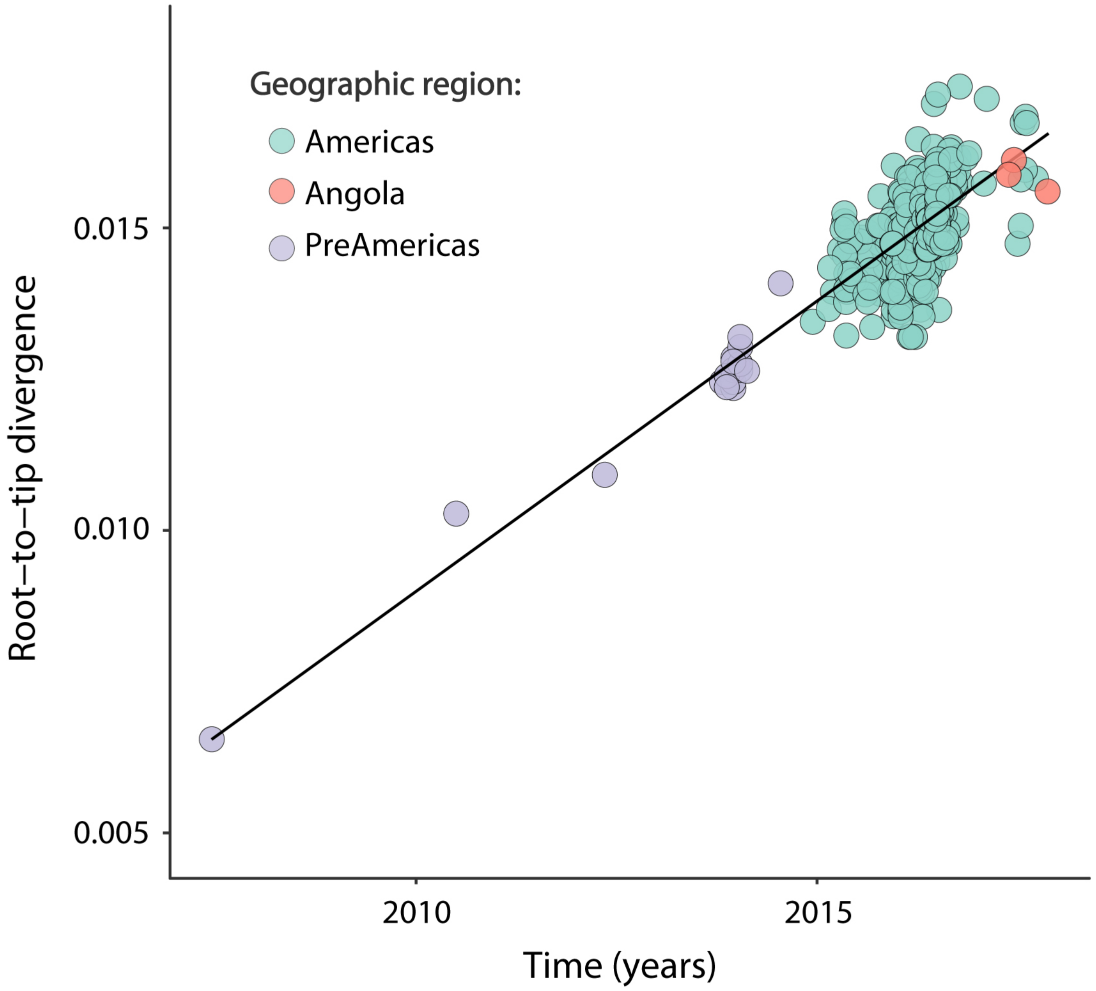


**Supplementary Figure 2. Root to tip regression of sequence sampling date and genetic divergence and maximum likelihood phylogeny.** Sequences are coloured by sampling location.

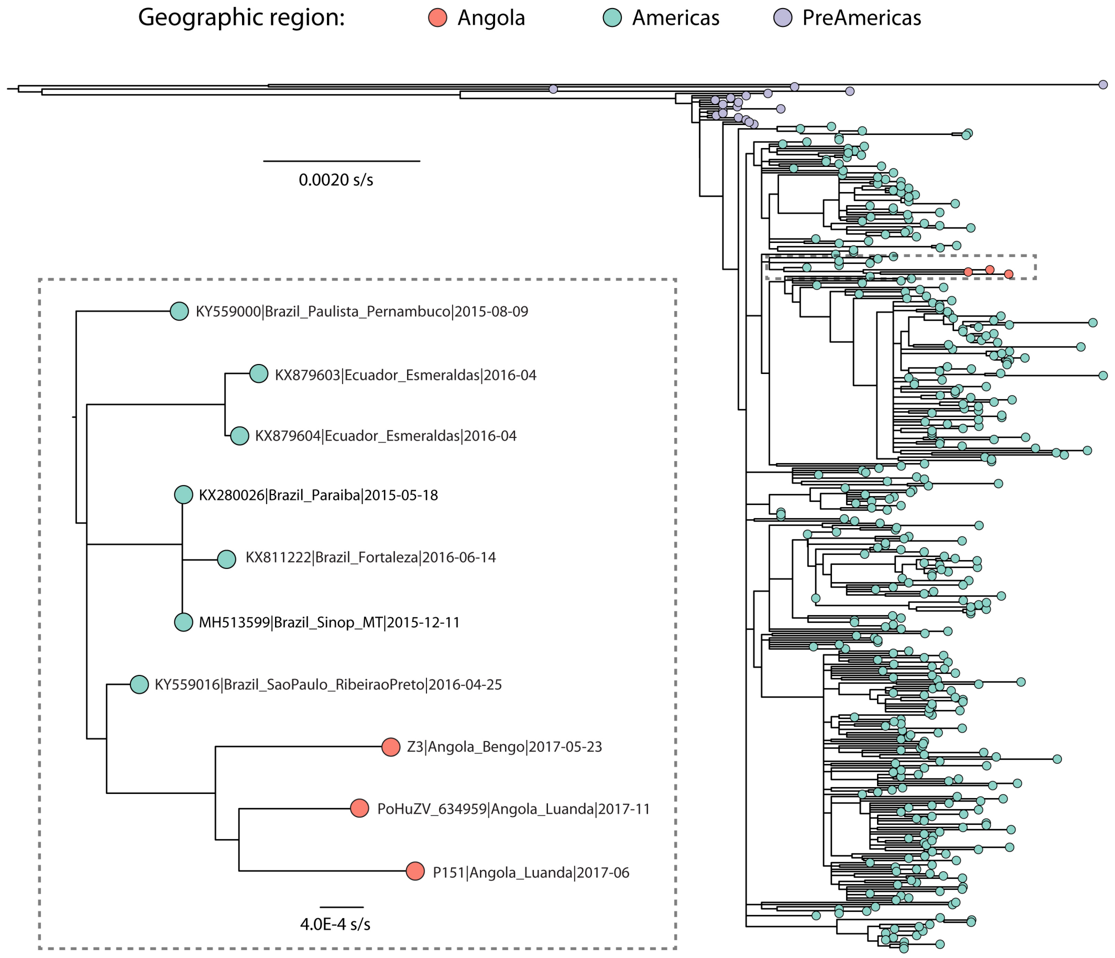


**Supplementary Figure 3. Maximum likelihood phylogeny of ZIKV.** The tree was estimated using 393 complete and partial ZIKV genomes (see Methods for details). The clade containing the Angola sequences is expanded on the bottom left. Sequences are coloured by sampling location (as in Supplementary Figure 2).

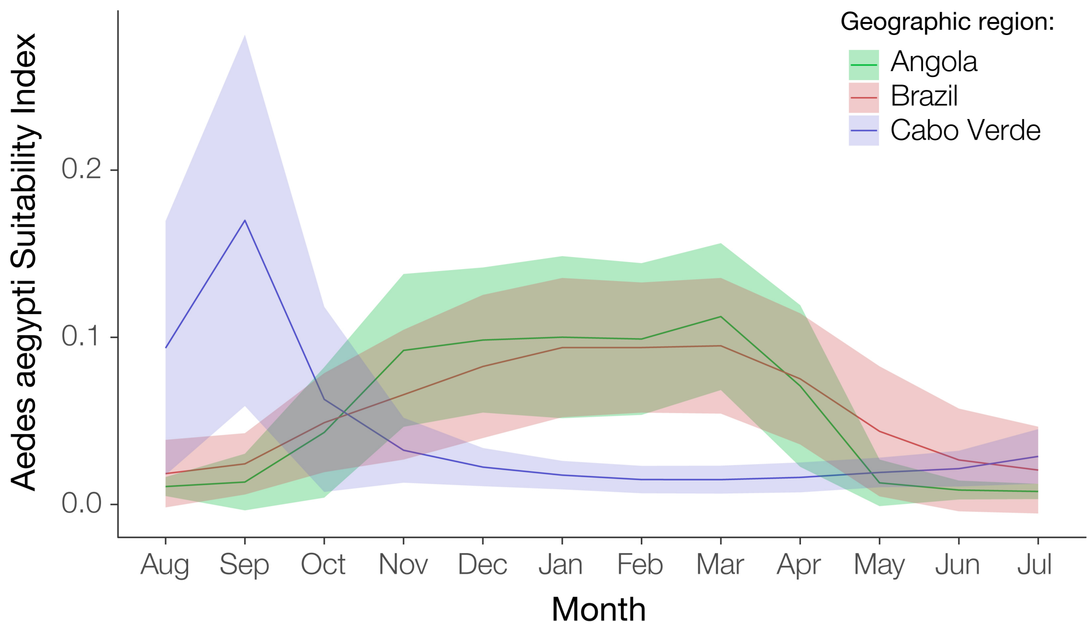


**Supplementary Figure 4. Seasonal climatic suitability for *Aedes aegypti*.** Solid lines represent the mean climatic suitability in that month and shaded areas show 68% confidence intervals. Data were obtained from [13]. Mean suitability per country was calculated using the R *raster* package (https://cran.r-project.org/web/packages/raster/index.html)

**Supplementary Table 1. Sequencing statistics for Angolan ZIKV genomes.** Note that a short 193bp fragment of the ZIKV NS1 gene from one patient (a microcephalic neonate born in Portugal) was previously reported (GenBank accession number MG742364).[14] As expected, this fragment exactly matches the relevant section of the larger genome sequence reported here, confirmed by a synonymous mutation observed in both sequences that is not seen in any other sampled ZIKV Asian lineage or African lineage genomes.

| **Sample (GenBank accession number)** | **Genomic coverage (%)** | **Mapped reads** | **Average depth** | **Bases covered >10x** | **Bases covered >25x** |
|---|---|---|---|---|---|
| Z3 (XXX) | 64·3 | 215916 | 3051 | 8910 | 8287 |
| P151 (XXX) | 80·8 | 17162 | 714 | 9236 | 8924 |
| PoHuZV/634959 (XXX) | 88·6 | 530192 | 3961 | 10249 | 10193 |

(GenBank accession numbers are pending and will be added prior to acceptance)
